# The Lineage-Specific Transcription Factor CDX2 Navigates Dynamic Chromatin to Control Distinct Stages of Intestine Development

**DOI:** 10.1101/425827

**Authors:** Namit Kumar, Yu-Hwai Tsai, Lei Chen, Anbo Zhou, Kushal K. Banerjee, Madhurima Saxena, Sha Huang, Jinchuan Xing, Ramesh A. Shivdasani, Jason R. Spence, Michael P. Verzi

**Affiliations:** Rutgers, the State University of New Jersey, Department of Genetics; Cancer Institute of New Jersey, and Human Genetics Institute of New Jersey, Piscataway, NJ, 08854, USA; Department of Internal Medicine; Department of Cell and Developmental Biology; Center for Organogenesis University of Michigan Medical School, Ann Arbor, MI 48109, USA; Department of Biomedical Engineering University of Michigan College of Engineering, Ann Arbor, MI 48109, USA; Department of Medical Oncology and Center for Functional Cancer Epigenetics, Dana-Farber Cancer Institute, Boston, MA 02215, USA; Department of Medicine, Brigham and Women’s Hospital and Harvard Medical School, Boston, MA 02215, USA; Harvard Stem Cell Institute, Cambridge, MA 02139, USA

**Keywords:** lineage-specifying, transcription factor, development, patterning, intestine, chromatin

## Abstract

Lineage-restricted transcription factors, such as the intestine-specifying factor CDX2, often have dual requirements across developmental time. Embryonic-loss of CDX2 triggers homeotic transformation of intestinal fate, while adult-onset *Cdx2*-loss compromises critical physiological functions but preserves intestinal identity. It is unclear how such diverse requirements are executed across the developmental continuum. Using primary and engineered human tissues, mouse genetics, and a multi-omics approach, we demonstrate that divergent CDX2 loss-of-function phenotypes in embryonic versus adult intestines correspond to divergent CDX2 chromatin-binding profiles in embryonic versus adult stages. CDX2 binds and activates distinct target genes in developing versus adult mouse and human intestinal cells. We find that temporal shifts in chromatin accessibility correspond to these context-specific CDX2 activities. Thus, CDX2 is not sufficient to activate a mature intestinal program, but rather, CDX2 responds to its environment, targeting stage-specific genes to contribute to either intestinal patterning or maturity. This study provides insights into the mechanisms through which lineage-specific regulatory factors achieve divergent functions over developmental time.

## Introduction

Lineage-specifying transcription factors, sometimes called “master regulators”, are required for a cellular lineage to develop in embryos and their loss leads to a selective failure to develop that lineage. Examples include MYOD, which establishes muscle development and induces muscle trans-differentiation in fibroblasts (Davis et al., 1987; Tapscott et al., 1988), or PDX1, which is essential for pancreas development (Jonsson et al., 1994; Offield et al., 1996). Study of lineage-specifying transcription factors is informative during early development, when tissues are specified; however, the same factors often have additional roles in subsequent tissue maturation, organogenesis, and adult homeostasis.

The homeodomain transcription factor, CDX2, is expressed from the onset of intestinal development through adult life. In mice, early deletion of *Cdx2* in the visceral endoderm precludes formation of the intestinal epithelium, which is replaced by an esophagus-like homeotic conversion (Gao et al., 2009). When CDX2 is lost early in the developing mouse endoderm, a striking patterning deficit follows, best illustrated by homeotic conversion of intestinal to esophageal epithelium (Gao, 2009), but also observed upon spontaneous loss of heterozygousity of CDX2 leading to harmatomas in the colon with features of esophageal squamous epithelium (Chawengsaksophak et al., 1997; Tamai et al., 1999), and expression of certain gastric markers in the intestinal epithelium upon early embryonic CDX2 inactivation using the Villin-Cre model (Grainger et al., 2010). In striking contrast to these findings, deletion of CDX2 at later stages in fetal and adult life compromises intestinal function, but not tissue identity (Verzi et al., 2010; Verzi et al., 2011), although tissue identity can become permissive when intestinal stem cells lacking CDX2 are isolated and subsequently cultured in a cocktail of factors conducive to gastric cell culture (Simmini et al., 2014), or upon prolonged loss of CDX2 in a subset of tissues (Hryniuk et al., 2012; Stringer et al., 2012).

The distinct consequences of CDX2 loss at different developmental stages prompts questions about the mechanistic basis of lineage-specifying transcription factor activities: Do these factors bind their chromatin targets from the outset or do their transcriptional targets differ along the developmental continuum? How do they interact with chromatin across developmental time? Is expression of a lineage-specific transcription factor sufficient to activate appropriate target genes, even in the absence of extracellular signals? Additionally, while many insights have been garnered from studies in mice, CDX2 function in human tissue specification remains untested.

Here, we employ mouse models and human pluripotent stem cell-derived models, coupled with investigation of chromatin accessibility, to show that CDX2 binds distinct chromatin sites in embryonic and adult intestines; this distinction is conserved in mice and humans. We find that CDX2 is incapable of instructing major shifts in the chromatin to direct its own binding. Rather, it follows the transitions in chromatin landscape that occur in the course of intestine development, and importantly, sustains that evolution. Our identification of distinct binding sites and regulatory functions during tissue specification and in adults provides one explanation for why intestinal identity in the absence of CDX2 is compromised only during a defined developmental window. These findings support a model in which lineage-specifying transcription factors operate in context-dependent roles, shaped by a dynamic developmental chromatin landscape.

## Results

### CDX2 is required for human intestinal development

In mouse embryos with conditional deletion of *Cdx2* in the developing endoderm, the intestinal epithelium exhibits morphologic and molecular characteristics of foregut identity. Esophageal, gastric corpus, or gastric antral identities are observed, depending on the timing of CDX2 loss, with earlier onset of *Cdx2* inactivation producing more anterior (rostral) identity shifts (Gao et al., 2009; Grainger et al., 2010). It is unknown whether CDX2 has a conserved role in human intestine development, and new methods to direct differentiation of human pluripotent stem cells (hPSCs) allow us to ask this question (D’Amour et al., 2005; Finkbeiner et al., 2015; McCracken et al., 2011; Spence et al., 2011).

CDX2 expression is first observed upon specification of the hindgut by WNT and FGF signaling (Wells and Melton, 2000). Immunoblotting confirmed absence of CDX2 in human embryonic stem cells (hESC) or definitive endoderm, but combined activation of WNT (using the GSK3β inhibitor, CHIR99021, to stabilize β-CATENIN) and FGF (using recombinant FGF4) signaling triggered robust CDX2 expression, which was sustained in 3-dimensional human intestinal organoid (HIO) cultures months later (Fig. 1A-C). To determine if *CDX2* is required to induce the intestinal lineage, we used Cas9/CRISPR to engineer *CDX2*-null hESC in the H9 line and differentiated the cells into HIOs. Sequencing confirmed mutation of *CDX2*, and CDX2 immunoreactivity was lost in mutant cells directed toward intestine differentiation (Fig. 1D and S1). Whereas control hESC gave rise to tissues that expressed CDX2 (Fig. 1D) and the intestine-restricted markers *AMY2B, MUC2*, and *DPP4* (Fig. 1E), *CDX2*-null hESCs failed to develop tissues with these characteristic markers. Instead, they gave rise to organoids expressing transcripts and proteins of CDX2-negative foregut lineages, including stomach, esophagus, pancreas, and gall bladder (Fig. 1D-E). These findings indicate a conserved role for CDX2 in initiating human intestine development and underscore the importance of defining the mechanisms through which CDX2 imparts intestinal fate.

**Figure 1.**
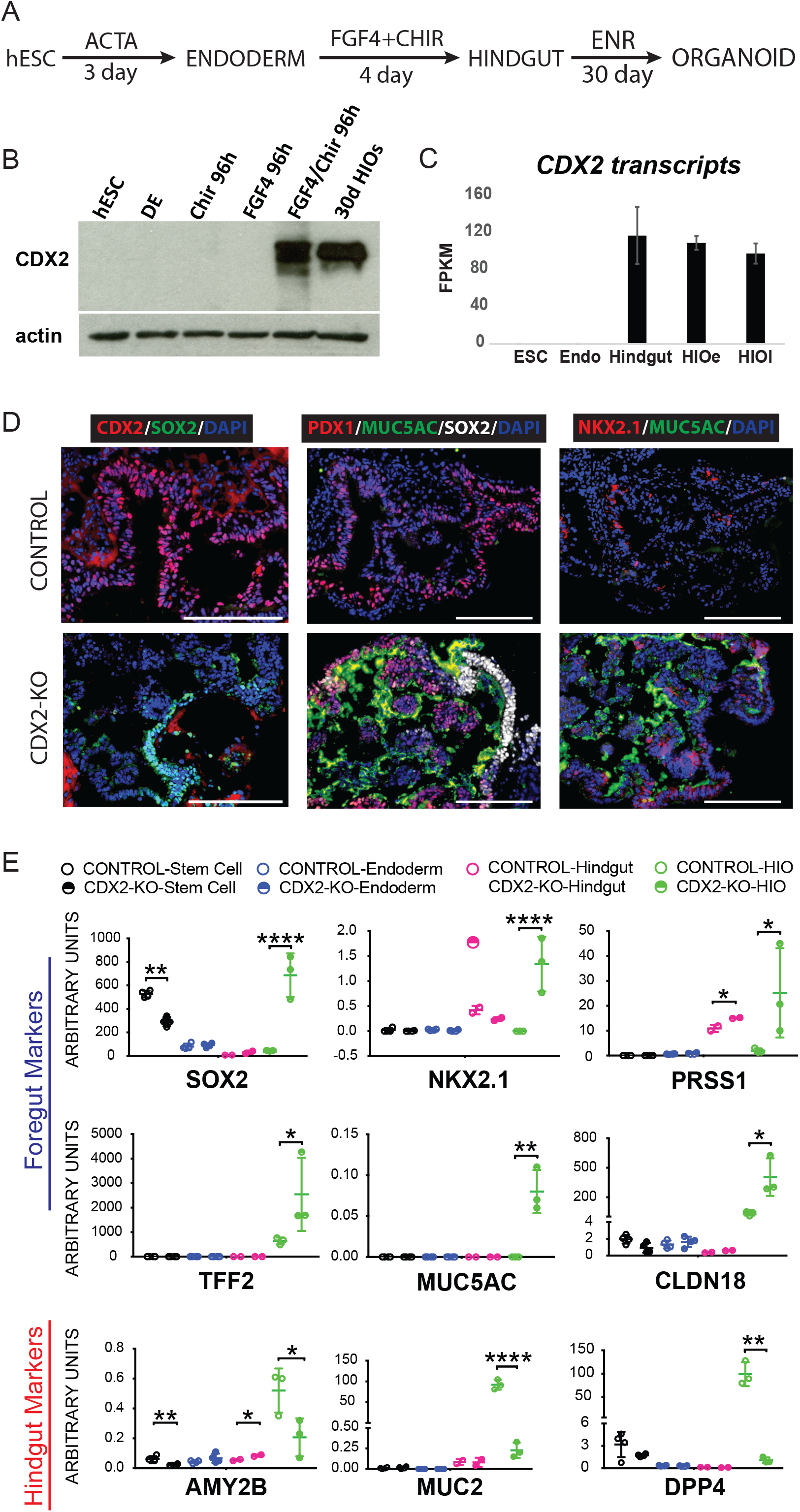
CDX2 is required for human intestinal development. A) Differentiation schema used to differentiate hES cells into Human Intestinal Organoids (HIOs). B) Immunoblot and C) RNAseq (E-MTAB-4168, (Tsai et al., 2017)) show that CDX2 expression is only induced robustly after differentiation of ES cells into endoderm with Activin and then induction of hindgut specification via treatment with FGF4 (500ng/ml) and the WNT agonist CHIR99021 (2µM). HIOe = HIO early, pre-transplant; HIOl = HIO late, after transplant into mouse kidney capsule for further intestinal maturation. D-E) While Human Intestinal Organoids (HIO) result from this differentiation protocol, ES cells lacking CDX2 due to Crispr targeting (Fig. S1) do not give rise to HIOs, and instead give rise to organoids expressing genes characteristic of foregut lineages, as seen by immunofluorescent staining (D, representative of at least 3 independent experiments) and qRT-PCR (E) on organoids collected after 30 days in ENR (EGF, Noggin, RSPO) culture conditions. (Unpaired t-test). Scale bars = 200µm.

### CDX2 regulates distinct transcriptional targets across human intestinal development

To determine these mechanisms, we performed ChIP-seq on biological replicates (Fig. S2A) of intestine-specified endoderm (herein referred to as ‘Hindgut’, as defined in Fig. 1A) and on primary adult human duodenal epithelium (herein referred to as ‘Adult’). We used MACS (Zhang et al., 2008) to identify ChIP-seq binding sites and *k*-means clustering to define genomic regions with differential CDX2 occupancy (Fig. 2A-B). Ontology analysis of genes linked to CDX2 binding sites revealed that Hindgut-enriched CDX2 binding occurred near genes that were enriched for functions in development, patterning, and morphogenesis, whereas genes near Adult-enriched CDX2 binding sites were associated with adult intestinal physiological functions (Fig. 2C). DNA sequence motifs enriched at Hindgut- and Adult-enriched regions were also distinct, suggesting that CDX2 works in different regulatory complexes in embryos and adults. The CDX2 motif was strongly enriched in both groups of binding regions, but whereas Hindgut-enriched sites were enriched in NANOG and SOX15 motifs, consistent with a role for SOX factors supporting endoderm development (Kanai-Azuma et al., 2002), Adult-enriched regions were most enriched in HNF4A motifs (Fig. 2D, Table S1), consistent with the function of HNF4 in the mature intestine (Babeu et al., 2009; Cattin et al., 2009; San Roman et al., 2015; Stegmann et al., 2006). Examples of Hindgut-enriched target genes included the *HOXB* cluster genes (Fig. 2E, S2B) known for their roles in tissue patterning and axial elongation (Young et al., 2009), whereas adult-enriched target genes included the enterocyte differentiation marker *ALPI* and the differentiation-promoting transcription factor gene *HNF4A* (Fig. 2E, S2C). These results reveal that human CDX2 binds distinct genomic regions in a stage-specific manner and regulates stage-specific genes.

**Figure 2.**
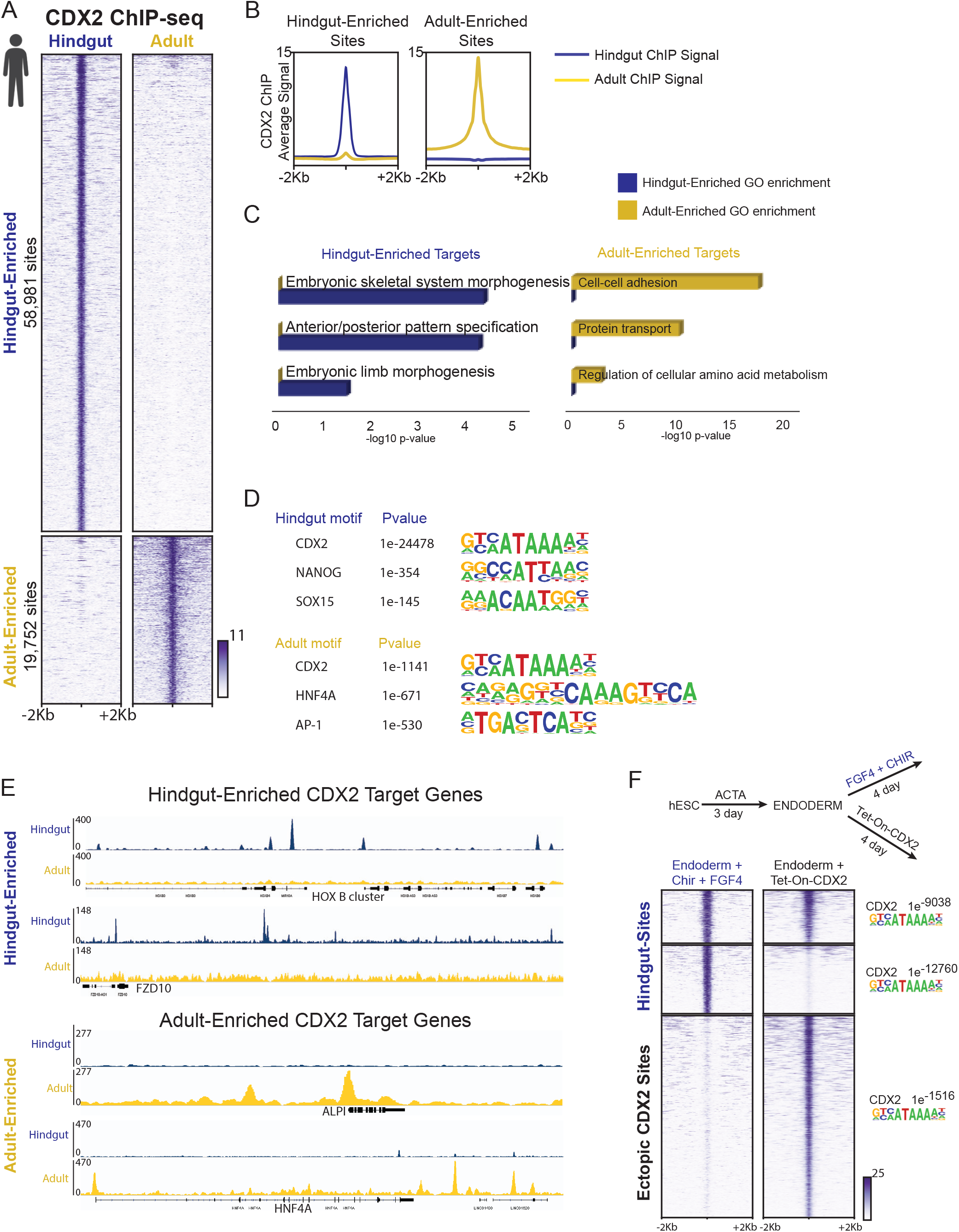
CDX2 has distinct transcriptional targets in the developing hindgut versus the adult intestine. A-B) CDX2 ChIP-seq performed in *in vitro* differentiated hindgut (4 day FGF4+CHIR treated endoderm, Fig. 1A) and in primary adult human duodenum epithelium show distinct binding profiles, as defined by *k*-means clustering (A) and compared using composite signal traces (B). (2 biological replicates each, Figure S2A). C) Distinct binding sites correspond to different sets of nearby target genes (within 5 kb of binding sites), as indicated by GO analysis using DAVID. D) CDX2 binding motifs are present in both categories of binding regions, but other transcription factor binding motifs are differentially enriched in the hindgut-enriched or adult-enriched CDX2-binding regions, defined using HOMER. E) Examples of stage-specific CDX2 binding sites are shown using IGV. F) Ectopic expression of CDX2 in endoderm, using a Dox-inducible hES cell line (Endoderm + 4 days DOX treatment), is not sufficient to recapitulate CDX2 binding to the majority of its normal hindgut target sites (Endoderm + 4 days CHIR + FGF). Approximately 52% of CDX2 hindgut sites (MACS p < 1e^-10^) were not detected by MACS in the Dox-inducible condition, even at a lower peak-calling stringency (p < 1e^-3^). Lack of binding cannot be attributed to differences in CDX2 binding motifs, which were robust in each category of binding site (HOMER). Also see Figure S2 and Table S2.

Both WNT and FGF signals are required to drive robust differentiation of human endoderm into hindgut (Spence et al., 2011) and activation of these signals induces CDX2 expression (Fig. 1A). It is unclear whether CDX2 expression is sufficient for its binding at hindgut regulatory regions or whether CDX2 also requires other sequelae of dual signaling pathway activation to arrive at its hindgut binding regions. To address this question, we engineered hESC for doxycycline-inducible CDX2 expression. ChIP-Seq after forced CDX2 expression in endoderm cells yielded more than twice as many binding sites as were present in CHIR- and FGF4-treated endoderm (135,194 sites compared to 58,981 sites, Fig. 2F), and both groups of regions were enriched in the CDX2 consensus motif (Fig. 2F, Table S2), consistent with *bona fide* CDX2 binding to these regions. Surprisingly, despite robust expression (Fig. S2E) and occupancy of significantly more genomic sites, ectopic CDX2 failed to bind more than half of the Hindgut-enriched targets occupied in the presence of CHIR and FGF4 (Fig. 2F), despite robust enrichment of consensus CDX2 binding sequences at these regions (HOMER, p = 1e^-12760^, Fig. 2F, Table S2). Thus, while CDX2 expression is necessary for specification of human intestine (Fig. 1C-D), CDX2 cannot reach its full complement of hindgut chromatin targets in the absence of WNT and FGF signaling cues.

### Dynamic CDX2 activity across intestinal development is conserved in mice

To refine and corroborate this model of CDX2 binding and activity *in vivo*, we turned to mice. First we identified CDX2 binding sites in intestinal cells isolated at embryonic days (E) 13.5, 16.5 and 17.5, and from adult intestines. *k* -means clustering identified 4 major groups: 4,123 genomic regions were bound more robustly in embryos, 12,551 sites were more specific to the adult (Fig. 3A-B), and the remaining sites were bound similarly at all stages (1,791 sites with robust CDX2 binding and 15,368 sites with modest signals – clusters 1 and 4 in Fig. S3A and Table S3). CDX2 showed a notable transition late in gestation: binding at embryo-enriched sites declined by E16.5, as occupancy of adult-enriched regions became evident (Figure 3A). To test stage-specific binding functions in gene regulation, we purified control and CDX2-depleted epithelium after embryonic knockout (*Shh-Cre;Cdx2^f/f^*, collected at E12.5) or adult knockout (*Villin-Cre^ERT2^;Cdx2^f/f^*) and used RNA-seq to identify CDX2-dependent transcripts. Consistent with CDX2’s activity as a direct transcriptional activator, the genes nearby CDX2 binding sites showed reduced expression upon CDX2 loss at the corresponding developmental stages (Fig. S3B). Developmental stage-specific CDX2 binding regions also correlated with stage-specific gene expression, as genes linked to adult-enriched binding sites were expressed more robustly in adult than in the developing intestine, and the converse was true for embryo-enriched binding targets (Fig. 3C). Thus, stage-specific CDX2 binding predicts nearby gene expression in each context.

**Figure 3.**
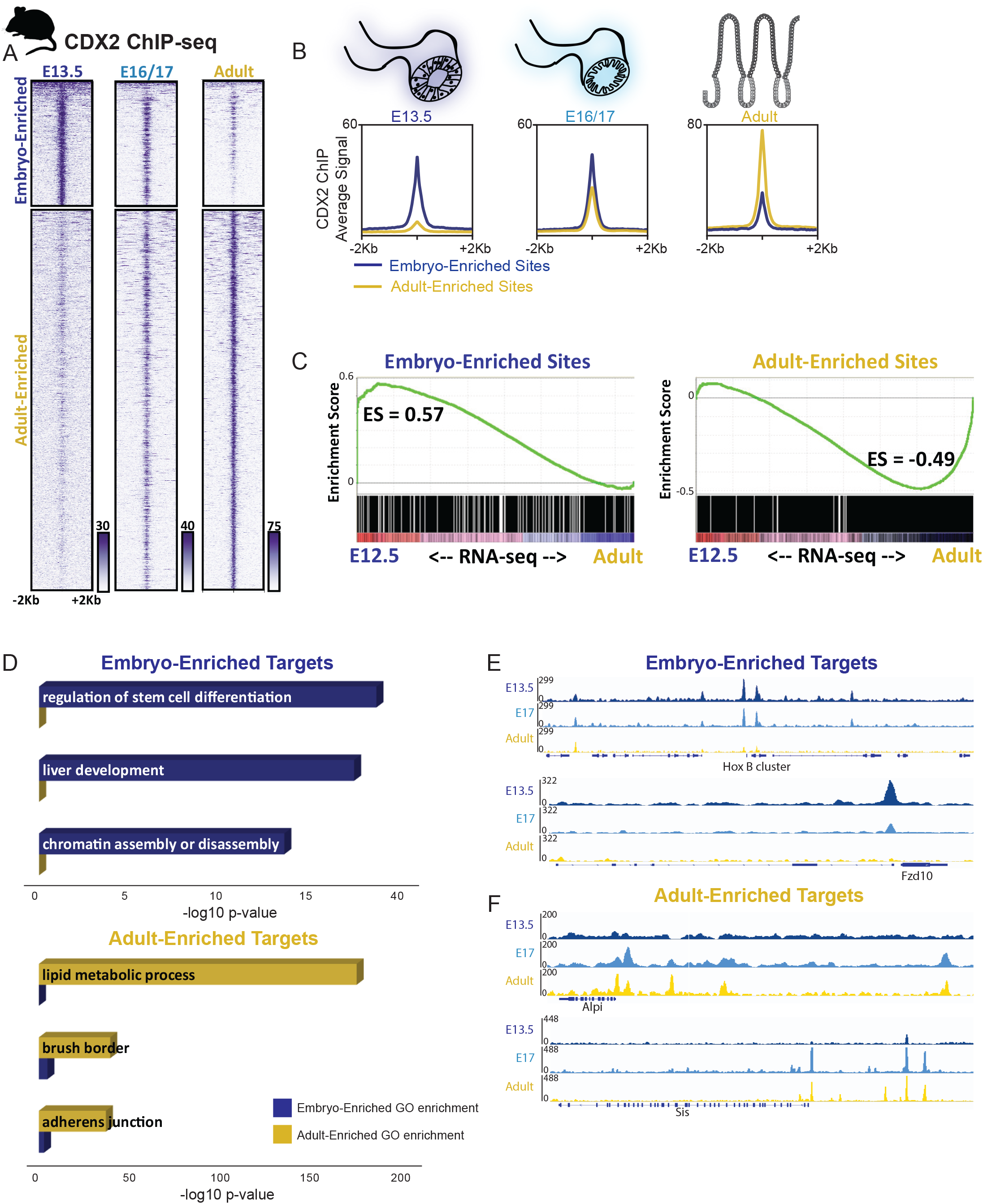
Temporal-specific shifts in CDX2 binding is conserved between mice and humans, and corresponds to temporal shifts in intestinal gene expression. A) CDX2 ChIP-seq performed at the indicated stages of intestinal development on isolated epithelium show distinct, stage-dependent binding profiles (2 biological replicates each, Figure S3). The heatmap depicts CDX2 ChIP-seq intensity at each of the regions defined as Embryo or Adult-enriched, and in (B) the average ChIP-seq signals at these binding regions is plotted. C) Embryo-enriched and Adult-enriched binding sites correspond to different sets of nearby target genes, which show stage-specific gene expression. RNA-seq was conducted on E12.5 or adult intestinal epithelium and genes linked to Embryo-enriched or Adult-enriched CDX2 binding (within 5kb) were analyzed for their distribution along the E12.5-to-adult expression continuum using GSEA analysis. CDX2-bound genes are also dependent upon CDX2 for expression, as revealed by RNA-seq analysis in *Cdx2* knockout tissues (Figure S3B), and have distinct gene ontologies, corresponding to embryo-specific or adult-specific functions (D). Representative examples of Embryo-enriched (E) or Adult-enriched (F) CDX2 binding sites are shown using IGV. Also see Figure S3.

Transcription factor motifs enriched at stage-specific enhancers also suggest dynamic regulatory activity. While CDX2 binding motifs were enriched in both adult- and embryo-enriched binding sites, other motifs were selectively enriched. For example, embryo-enriched CDX2 binding was associated with HOX and FOXA motifs, whereas Adult-enriched binding sites were associated with HNF4, FRA1, and KLF5 motifs (Fig. S3C, Table S3). This divergence points to different transcriptional regulatory networks in the embryonic gut, where developmental competence and tissue patterning predominate (Wang et al., 2015) and in the adult intestine, where cell identity is fixed and CDX2 and partner factors control physiologic functions (Verzi et al., 2010; Verzi et al., 2011). Indeed, ontology analysis indicated that genes near embryo-enriched binding sites associate with developmental processes such as liver development and regulation of stem cell differentiation, whereas genes near Adult-enriched regions associate with mature intestinal functions, including lipid metabolism and brush border (Fig. 3D). Condition-specific binding at developmental genes (*HoxB, Fzd10*) or those expressed exclusively in the mature tissue (*Alpi, Sis*) illustrates these findings (Fig. 3E-F, S3D-E). Together, these results reveal that CDX2 has distinct genomic targets at different developmental stages, regulates target genes in a stage-dependent manner, and likely partners with different factors in each context to accomplish stage-dependent gene regulation.

### Temporal dissection of *Cdx2* requirements for intestinal identity

We next sought to correlate our observed changes in CDX2’s genomic occupancy with observations of homeotic conversion phenotypes upon CDX2 loss. CDX2 loss triggers profound homeotic transformation of the intestine to esophagus when it occurs at E8.5 (*FoxA3-Cre*, (Gao et al., 2009). When Cdx2 is deleted at E13.5, however certain stomach-specific genes are ectopically expressed without overt squamous cell differentiation (Grainger et al., 2010). We examined the timing of CDX2 dependency more closely, with additional time points of CDX2 inactivation. Mouse endoderm engineered to delete *Cdx2* at ~E9.5 using the *Shh-Cre* driver (Harfe et al., 2004) exhibited stomach characteristics anteriorly in the jejunum (ATP4B and foveolar PAS staining) and stratified squamous esophageal features posteriorly (Fig. 4A, Fig. S4A). Thus loss of CDX2 ~E9.5 results in a homeotic change of intermediate phenotypic severity compared to when CDX2 is lost earlier E8.5 (Gao et al., 2009), or later E13.5 (Grainger et al., 2010). In contrast, we observed no evidence of transformation when we inactivated *Cdx2* at later embryonic stages by tamoxifen treatment of dams carrying *Villin-Cre^ERT2^*; *Cdx2^f/f^* embryos at E13.5 or E15.5 (Fig. 4A). Thus these late stage embryos, like the adult epithelium upon CDX2-loss (Verzi et al., 2011), retain intestinal identity. A summary of our findings on the consequences of CDX2 inactivation in the developing intestine, along with those previously published, indicates a critical window prior to E13.5 in which intestinal identity is plastic; after this timepoint, intestinal identity is stabilized and less-susceptible to transformation to anterior structures (Fig. 4B). These temporal phenotypes roughly correspond to the dynamic CDX2 binding patterns we observed in ChIP-seq assays of early versus late embryonic intestinal epithelia (Figure 3A) and are consistent with observations that CDX2 target genes partition into functions associated more with developmental process (Embryo-enriched sites) versus mature intestinal physiological functions (Adult-enriched sites, Figure 3D).

**Figure 4.**
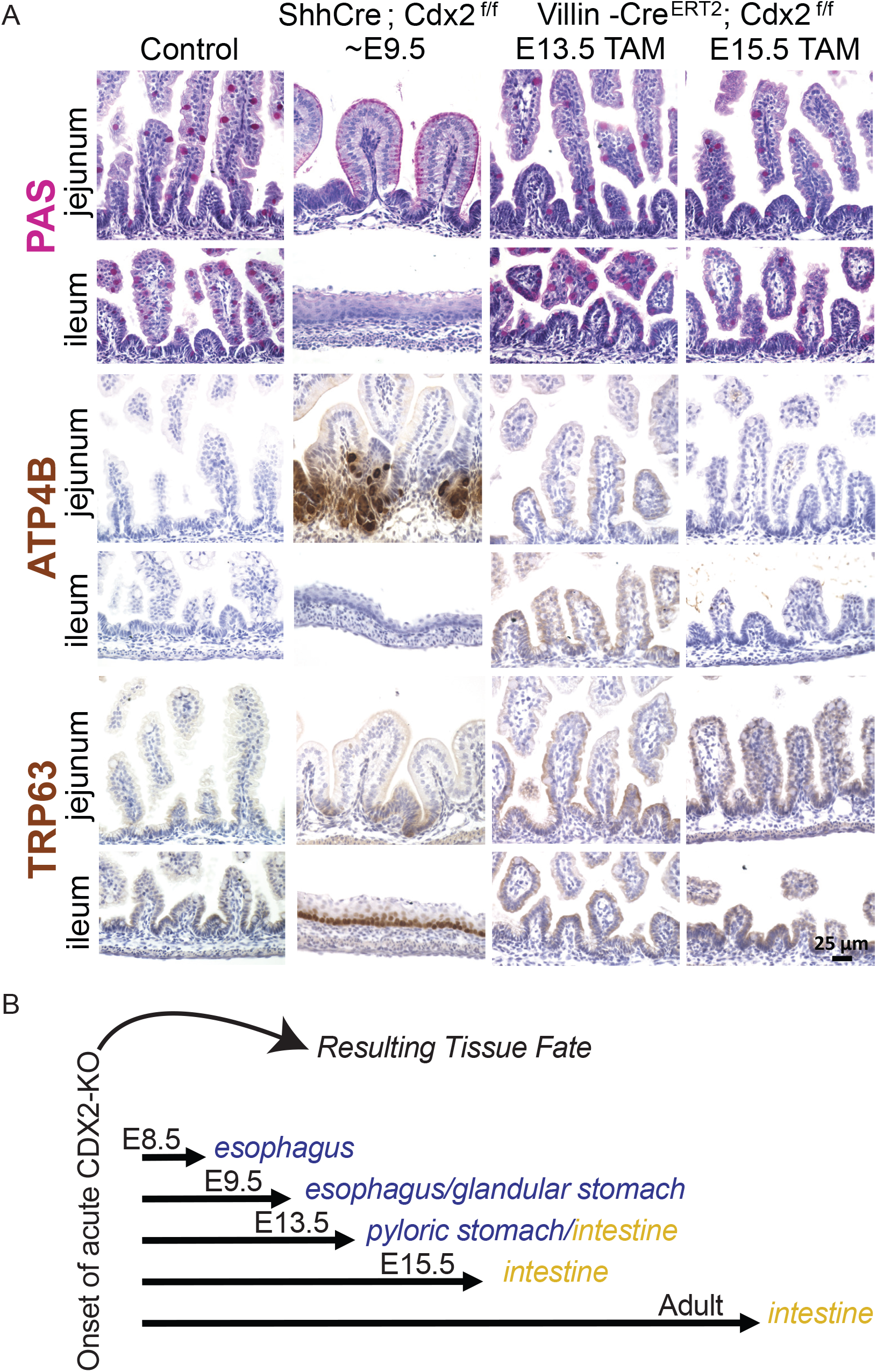
Developmental stage dictates the consequences of CDX2 loss. A) Analysis of regional tissue identity on samples from control, *Shh-Cre; CDX2^f/f^* or *Villin-Cre^ERT2^; CDX2^f/f^* embryos, treated with tamoxifen to induce CDX2 loss as indicated, and collected at E18.5 from the indicated intestinal region. PAS stain marks mucosubstances, including intestinal Goblet cells, which are identified by a punctate staining and are characteristic in the intestinal epithelium (Control or Villin-Cre^ERT2^), whereas broad PAS staining at the apical cell surface is indicative of gastric foveolar cells, as observed in the jejunum of *Shh-Cre*; *Cdx2^f/f^* embryos. TRP63 marks esophageal cells and is not typically found in the intestine, but appears ectopically in the ileum of *Shh-Cre*; *Cdx2^f/f^* embryos, but not in control embryos, or embryos induced to delete *Cdx2* in the later stages. The parietal cell marker, ATP4B, is typically expressed in the glandular stomach, but is ectopically expressed in the CDX2-negative jejunum of *Shh-Cre*; *Cdx2^f/f^* embryos, but not in embryos deficient in intestinal CDX2 at the later stages tested. B) Summary of CDX2 loss-of-function phenotypes from this (A) and previous studies. Temporal-specific CDX2 loss at stages early than E12.5 induces homeotic transformations, as observed by Gao and colleagues using the *Foxa3-Cre* driver (~E8.5), in this study using the *Shh-Cre* driver (~E9.5), or at ~E13.5 using the *VIllin-Cre* driver (Grainger et al., 2010). However, acute loss of CDX2 at later developmental stages (>E13.5) and in the adult compromises intestinal digestive functions but intestinal identity is preserved (*Villin-Cre^ERT2^* with tamoxifen treatment at E13.5, E15.5, or adult), as documented in these studies in the embryo (Gao et al., 2009; Grainger et al., 2010), and previously in the adult (Verzi et al., 2011).

### CDX2 follows, then stabilizes, a developmentally dynamic chromatin landscape

CDX2 expression is nearly constant from the onset of intestine development through adult life in both humans (Fig. 1A-B) and mice (Fig. 5A-B). To resolve why CDX2 could function in different capacities despite relatively consistent levels of expression (Figs. 2-3), we examined the possibility that a dynamic chromatin environment could underlie stage-specific CDX2 genomic binding. Chromatin accessibility was measured at all embryonic and adult binding sites using ATAC-seq at E11.5, E14.5, E16.5, E18.5, P1, and Adult. We observed a clear pattern at CDX2-bound regions, with embryo-enriched sites losing accessible chromatin after E14.5, while adult-enriched sites gained chromatin access thereafter (Fig. 5C-D). Four features are notable in this regard. First, 74% of genomic regions with strong adult-enriched CDX2 binding showed poor accessibility in early embryos, lacking ATAC signals at E11.5 (*P* <10^-5^; Fig. 5C). Second, open chromatin emerged at these sites in adults (Fig. 5C), where CDX2 binding was clearly absent in the embryo (Figure 3A); the converse was apparent for the embryo-enriched CDX2 binding sites. Third, chromatin dynamics were similar for the active histone mark H3K27ac (Kazakevych et al., 2017), with stage-specific enhancer marks corresponding to stage-specific CDX2 binding regions (Fig. S5A-B). Fourth, stage-specific binding in mouse endoderm resembled the profound shift in binding between human ESC-derived gut endoderm and adult human intestine (Fig. 2A). Thus, despite constant expression across time, CDX2 binding closely follows a temporal wave of accessible chromatin from embryo-enriched to adult-enriched sites. Examples of these findings are illustrated at genes selectively expressed in developing or adult intestine (Fig. 5E-F).

**Figure 5.**
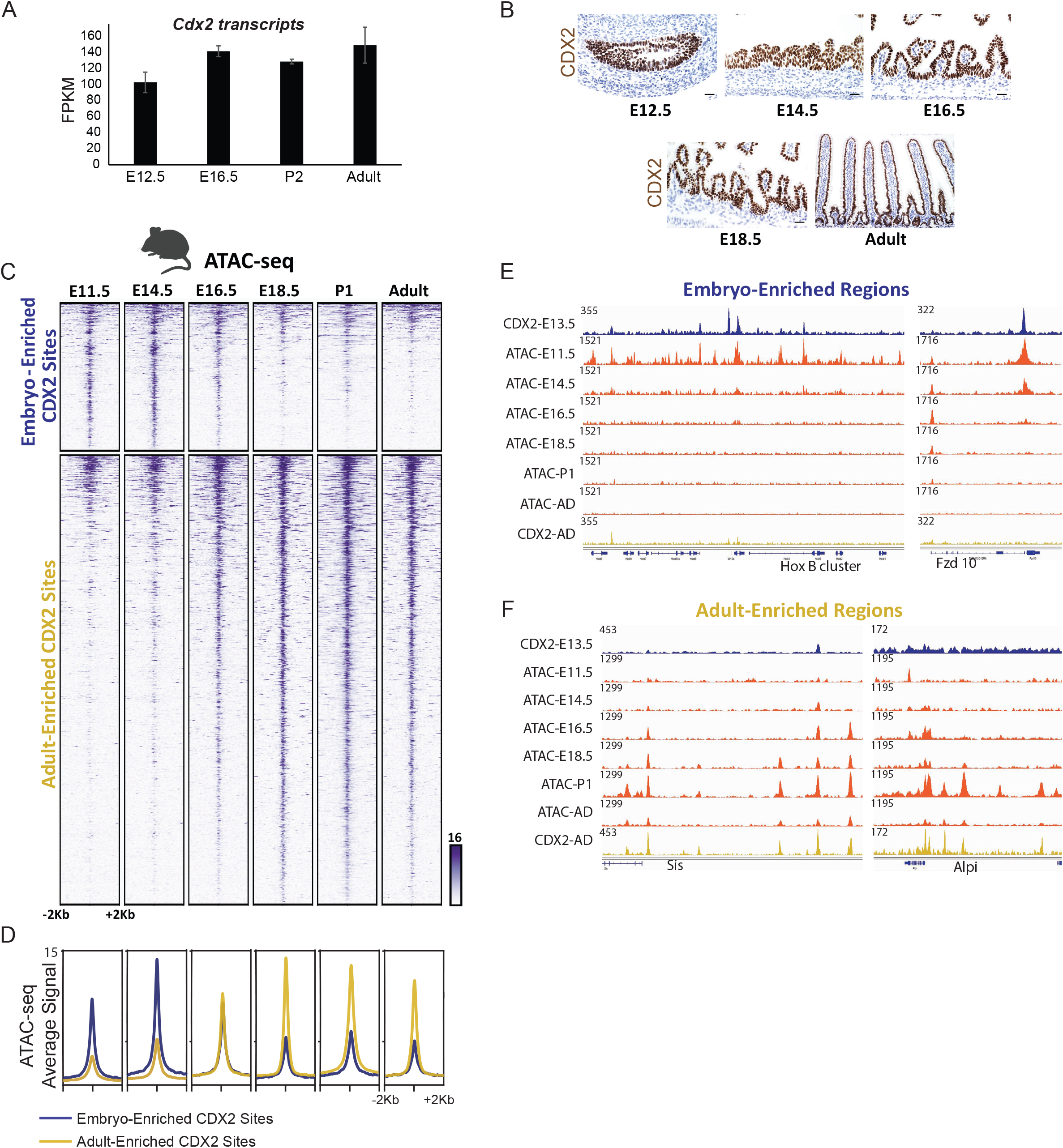
CDX2 temporally-enriched binding patterns follow a temporally dynamic chromatin landscape that requires CDX2 for maintenance. A) CDX2 transcripts (from RNA-seq on isolated epithelium) and B) protein levels (via immunostaining) are consistent across mouse developmental time, yet CDX2 occupies distinct sets of genomic regions at different developmental stages (Fig. 3). CDX2 binding chromatin binding shifts may be due to a shifting chromatin landscape across developmental time, as indicated by the ATAC-seq assay for accessible chromatin (C), shown at CDX2’s condition-enriched binding regions (defined in Fig. 3A). D) Average ATAC-seq signal is robust at embryo-enriched CDX2 binding sites prior to E16.5, after which chromatin accessibility at these regions diminishes. The converse is true for adult-enriched CDX2 binding sites. (E-F) Specific examples of condition-enriched chromatin accessibility at CDX2 binding sites. CDX2 binding regions are also dependent upon CDX2-binding to remain accessible to the ATAC-seq assay, as seen in CDX2 knockout epithelium (Figure S5).

The contemporaneous transitions we observed after E14.5 in chromatin accessibility (Fig. 5C-D), CDX2 binding (Figure 3A), and histone marks (Figure S5A-B) coincide roughly with the temporal restriction of *Cdx2*-null phenotypes (Fig. 4A-B). Importantly, although CDX2 cannot access adult-enriched target sites in the developmental context (Fig. 2A & 3A), ATAC-seq in *Shh-Cre*;*CDX2^f/f^* embryos at E16.5 revealed diminished chromatin accessibility at the vast majority of regions occupied by CDX2 at that developmental time point (Fig. S5C-D). Reduced chromatin access was specific to CDX2-bound regions, as chromatin accessibility at all annotated promoters was unchanged (Fig. S5D-E). Together, these data reveal that although chromatin accessibility directs CDX2 binding during endoderm development, CDX2 is required to sustain access at regions associated with adult intestinal genes, consistent with CDX2 control of enhancer chromatin structures in mature mice (Verzi et al., 2013). Thus, CDX2 acts in distinct developmental contexts, constrained in part by temporal chromatin landscapes, to promote tissue specification in embryos and essential digestive functions in adults.

## Discussion

Pioneer factors help shape accessible chromatin (Iwafuchi-Doi and Zaret, 2016) and developmental signaling pathways and partner transcription factors prime chromatin structure to create a state of “developmental competence” (Wang et al., 2015). Our findings fit within this model, in that CDX2 is unable to bind condition-enriched binding sites (embryo or adult) outside of the proper developmental context. Our findings demonstrate that CDX2 is repurposed across the transitions in developmental competence: first CDX2 binds genes associated with developmental and specification functions, and subsequently CDX2 binds genes that promote the mature differentiated state in adults. The correlation between chromatin accessibility (Fig. 5), active histone modifications (Fig. S5), and CDX2 binding suggests that the chromatin environment is likely a critical factor in dictating condition-specific binding of CDX2. Our experiments with forced CDX2 expression in hESC-derived endoderm are similar to our findings in the mouse, with the observation CDX2 is not sufficient to access at many of its primary hindgut targets independently of the signaling environment. These findings are consistent with CDX2’s ability to pioneer certain binding sites in ES cells, while depending on accessible chromatin at others (Mahony et al., 2014). Alternatively, post-translational modifications, expression of transcriptional co-regulators, or other regulatory mechanisms likely function downstream of WNT and FGF signaling to create an environment in which CDX2 can access its hindgut target sites and impart intestinal identity. A similar set of regulatory mechanisms could govern the transition of CDX2 from Embryo-enriched binding sites to Adult-enriched binding sites (Figures 2-3) across critical developmental transitions (Figure 4B). CDX2 binding during intestinal specification is likely to rely upon pioneer factors, and motif analysis points to FoxA, a known pioneer factor (McPherson et al., 1993), as a likely candidate (Figure S3C). As yet unidentified chromatin regulators likely generate the corresponding competence for CDX2 occupancy and intestinal maturation as the intestinal chromatin landscape is altered in late gestation (Figure 5C). Our observations that CDX2 binding and chromatin access shift during development provide new insights into the molecular basis of intestinal maturation and indicate that mere expression of lineage regulators is insufficient to achieve desired tissue fates. Defining the factors that craft embryonic chromatin landscapes toward their mature forms will bolster efforts of regenerative medicine to accurately develop these tissues *in vitro*.

In models of Barrett’s esophagus, a predisposing condition of esophageal adenocarcinoma, CDX2 is unable to induce intestinal metaplasia simply by its lone ectopic expression in the foregut; pathologic changes are triggered only within the proper cellular context (Jiang et al., 2017; Silberg et al., 2002). This limitation is consistent with the mechanisms we report. CDX2 does not bind mouse adult-specific intestinal regions in early endoderm. Only after the epigenome is restructured late in gestation does CDX2 access adult-specific binding regions. Consistent with the observation that the intestinal chromatin landscape shifts late in development, loss of CDX2 in the adult mouse epithelium leads to a fetus-like transcriptome (Mustata et al., 2013), but not the early embryonic state permissive to homeotic transformation. CDX2 loss in early embryos permits such transformation in both mice and humans (Gao et al., 2009; Grainger et al., 2010), Figures 4A-B), likely because a distinct chromatin environment is present for a limited time. Defining whether and how chromatin accessibility is altered in instances of homeotic transformation upon prolonged loss of CDX2 in adult tissues (Simmini et al., 2014; Stringer et al., 2008) will be an important next step in understanding the mechanisms of lineage plasticity.

## Materials and Methods

### Mice

*Shh-cre* mice (Harfe et al., 2004) were purchased from Jackson Labs and male *Shh-cre; Cdx2f/+* were bred to *Cdx2 ^f/f^* female mice for inducing Cdx2 knockout (KO) in the developing intestine beginning at ~E9.5. Cre-negative embryos were used as littermate controls. *Villin-Cre^ERT2^* (el Marjou et al., 2004) and *CDX2^f/f^* mice (Verzi et al., 2010) were previously described. Wild-type CD1 mice were obtained from Charles River Laboratories. All mouse protocols and experiments were approved by Rutgers Institutional Animal Care and Use Committee.

Key features of the methodologies are noted below and expanded upon in the supplemental methods.

### Human pluripotent stem cell culture and differentiation

Human embryonic stem cells (line H9, WiCell Research Institute, NIH registration number 0062) were maintained and differentiated into endoderm, hindgut and intestinal organoids as published previously (Cruz-Acuna et al., 2017; Tsai et al., 2016; Tsai et al., 2017).

### Histology, immunofluorescence and immunohistochemistry

Human organoids derived from stem cells (Spence et al., 2011) for 30 days were processed for immunofluorescence using antibodies (CDX2, Bio-Genex, MU392-UC, 1:500; MUC5AC, Abcam, ab79082, 1:500; NKX2.1, ThermoFisher, 8G7G3, 1:50; PDX1, Epitomics Inc, 3470-1 1:300; SOX2, Santa Cruz, sc-27603, 1:100). Mouse immunostaining was performed with following antibodies: CDX2 1:200, Cell Signaling, 12306; ATP4B 1:200, MBL International Corp., D032-3; TRP63 1:500, Santa Cruz Biotech, sc-8343.

### Purification of Intestinal epithelial cells from mice

Single cells from dissected intestines were incubated with PE-conjugated anti-CD326 (EpCam clone G8.8, eBiosciences, 12-5791-81) and PE-stained cells were then incubated for ~30 minutes with magnetic conjugated anti-PE antibody (Miltenyi Biotec Anti-PE MicroBeads, 130-048-801) and isolated on a column (Miltenyi Biotec, MS Columns, 130-042-201) in a magnetic field to obtain magnetic antibody conjugated, EpCam positive epithelial cells.

### ChIP-seq

For embryonic CDX2 ChIP-seq (E13.5, 12 embryos pooled per replicate; E16.5, 3 embryos pooled per replicate; and E17.5, 2 embryos pooled per replicate) midguts (caudal stomach to rostral caecum) were collected cells were isolated, fixed, and sonicated. Chromatin was incubated overnight with CDX2 antibody (6µl, Bethyl A300-691A) ChIP DNA was used to prepare ChIP-seq libraries using Rubicon Genomics ThruPLEX DNA-seq Kit (R400427/R400428/R40048), fragment size selected using Pippin Prep and sequenced on Illumina HiSeq.

### ATAC-seq

25,000-50,000 isolated epithelial cells, from midguts (caudal stomach to rostral caecum, up to 2 embryos per sample) were used for ATAC-seq as described previously (Buenrostro et al., 2015).

### RNA-seq

For E12.5 RNA-seq, total RNA was extracted from epcam-enriched epithelial cells (as described above) from gut tissue caudal to the stomach (2-5 embryos pooled per sample) in trizol using RNeasy micro kit (QIAGEN, 74004). RNA-seq libraries were prepared using the SMARTer-Seq v4 Low Input mRNA library kit (Clonetech SMARTer 634888). Adult RNA data were curated from GSE70766 and GSE98724 (San Roman et al., 2015; Saxena et al., 2017).

### Computational analysis

Raw Sequencing reads (fastq) were quality checked using fastQC (v0.11.3), aligned to mouse (mm9) or human (hg19) genomes using Tophat2 (v2.1.0, RNA-seq) or bowtie2 (v2.2.6, ChIP- and ATAC-seq). For ChIP- and ATAC-seq, bam files were merged using samtools merge (0.1.19) for downstream analysis. Bigwigs were generated from bam files using deeptools bamCoverage v2.4.2, (Ramirez et al., 2014).

RNA-seq: Cxb files were generated using Cuffquant, normalized expression count tables were generated from cxb files using Cuffnorm (v2.2.1, library-norm-method quartile). Differential gene expression tables were computed using Cuffdiff (v2.2.1) and log2 fold change (FPKM+1) used for analysis.

For ChIP-seq and ATAC-seq: Peaks (bed file) were identified using MACS (1.4.1, Zhang et al., 2008). Two biological replicate ChIPs were performed for each condition, and replicates inspected by measuring Pearson correlation coefficients. Genes associated with ChIP peaks were identified using BETA minus and Enriched Motifs at ChIP peaks were identified using HOMER findMotifsGenome.pl (Heinz et al., 2010).

For comparing open chromatin regions (ATAC-seq) or differential CDX2 binding (ChIP-seq), bigwigs were generated (no normalization) and signal intensities were quantile normalized (v0.4.0, bin_size 50, log transformation) using Haystack (Pinello et al., 2018). *k* -means clustered heatmaps were generated from haystack normalized bigwigs using computeMatrix and plotHeatmap from deeptools package (v 2.4.2).

For ATAC-seq, two biological replicates from each developmental stage were performed, bam files were merged and bigwigs were quantile normalized to compare intensities of open chromatin signal. Embryonic and adult ChIP-seq data (Kazakevych et al., 2017) of the active chromatin marker (H3K27ac) was applied to Cdx2’s E13.5 and adult enriched genomic targets.

All RNA-seq, ATAC-seq, and ChIP-seq data generated in this study can be found at GEO with accession# GSE115314 (using reviewer code uvojyywmjnujtiz) and GSE115541 (reviewer code ozsryousxfexlqx).

## Supplemental Methods

### Human pluripotent stem cell culture and differentiation

hESCs were maintained in mTESR1 (Stem Cell Technologies) on hESC qualified Matrigel (Corning) in 6 well dishes. Prior to differentiation, hESCs were passaged as small clumps into 24 well Nunc™ Cell-Culture Treated multi-well plates with the Nunclon™ Delta surface treatment. When cultures were approximately 60% confluent, hESC media was replaced with endoderm differentiation media (RPMI1640 + 100ng/mL ActivinA (R&D Systems) for three days with supplemented with 0%, 0.2% and 2% HyClone FBS on subsequent days. Following endoderm differentiation, cells were exposed to hindgut differentiation media for 4-6 days, which consisted of RPMI1640 + 2% HyClone FBS + 2uM CHIR99021 + 500ng/mL FGF4 (made in house, as previously described (Leslie et al., 2015). Free-floating hindgut spheroids formed on the 4^th^-6^th^ days, and were subsequently placed in a droplet of Matrigel and were grown/expanded into human intestinal organoids (HIOs) for 28-35 days, in HIO media as previously described (Tsai et al., 2016), which consisted of Advanced DMEM/F12 media supplemented with 1X Pen/Strep, 1X B27 (Gibco, Life Technologies), 100ng/mL EGF, 5% NOGGIN conditioned media (Heijmans et al., 2013) and 5% R-SPONDIN2 conditioned media (Bell et al., 2008). For human patients, de-identified duodenal biopsies were collected with Rutgers IRB approval, cut longitudinally and epithelial cells were obtained by scrapping the luminal surface. Cells were washed twice with PBS, fixed with 1% formaldehyde (15min at 4°C + 30 minutes at RT), washed twice and cell pellets were flash frozen for subsequent chromatin pulldown.

### Histology, immunofluorescence and immunohistochemistry

Human organoids were collected and fixed in 4% PFA for 1 hour, followed by cryoprotection with 30% sucrose overnight. On the next day, tissues were embedded with OCT (Fisher, Tissue-Plus, 4585) and stored in -80°C. Tissues, were sectioned at 3 µm and immunofluorescent staining was carried out as standard methods by staining primary antibody overnight and 1 hour secondary antibody on the next day. For hindgut immunofluorescent staining, the monolayer of cells was fixed by 4% PFA for 20 minutes, followed by the same methods as organoids. Images were taken by Olympic microscope IX71.

For mouse immunostaining, intestinal tissue was dissected and fixed overnight in 4% paraformaldehyde at 4°C, washed with PBS, passed through increasing concentrations of an ethanol series and paraffin. 5-µm intestinal sections were cut from paraffin blocks, processed for immunostaining with the indicated primary antibodies, developed using the Vectastain ABC Kit (Vector Laboratories, PK6101) and counterstained with hematoxylin. A one-hour antigen retrieval step in 10mM sodium citrate solution under 15 psi pressure was used for all stains. Slides were incubated with primary antibody overnight at 4°C (CDX2 1:200, Cell Signaling, 12306; ATP4B 1:200, MBL International Corp., D032-3; TRP63 1:500, Santa Cruz Biotech, sc-8343). Periodic acid-Schiff (PAS) staining was conducted by treating slides with 0.5% periodic acid and staining with Schiff’s Reagent (Alfa Aesar J612171). Images were taken using a Retiga 1300CCD (Q-Imaging) camera and a Nikon Eclipse E800 microscope with the QC-Capture imaging software. Adjustments in contrast and sharpness, when made, were applied to complete figure panels in Adobe Photoshop.

### qRT-PCR

Briefly, RNA isolation was performed using MagMAXTM-96 total RNA isolation kit (Ambion, AM1830). SuperScript VILO cDNA synthesis kit (ThermoFisher, 11754250) was used to make cDNA from 200 ng RNA. cDNA levels were detected using QuantiTect SYBR Green (QIAGEN, 608056). Relative gene expression was plotted as Arbitrary Units, using the following formula: [2>(housekeeping gene Ct - gene Ct)] × 10,000.

### Generating CDX2 hESC knockout cell line using CRISPR-CAS9

The CDX2-KO was generated using the pCas-Guide-EF1a-GFP plasmid (Blue Heron Biotech LLC, Cat# GE100018) using the CDX2 sgRNA of targeting sequence (CCTCTCAGAGAGCCCCAGCGTGG) at the human CDX2 exon2 DNA binding domain. Plasmid was introduced into hES cells using electroporation (NEPA21 Electro-Kinetic Transfection System). 10µg total plasmid DNA was prepared and mixed with 1X10^6^ hES cells in transfection medium (OPIT-MEM, Gibco, 31985-062) to a total volume of 100 µL and added into an electroporation cuvette (Bulldog Bio, 12358-346). Cells plus DNA were electroporated using the following settings: poring pulse: 175 volt; transfer pulse: 20 volt. Immediately, the transfected cells were transferred to one well of a Matrigel coated 6-well plate with 1.5 mL mTeSR1 medium plus ROCK inhibitor (REAGENTS DIRECT, Y27632, 53-B85). After 24 hours, GFP positive cells were sorted (FACS) and plated in Matrigel coated 6-well plates with mTeSR1 medium plus ROCK inhibitor at low density. After 5-7 days culture, single colonies were manually picked and transferred to individual wells of a 24 well plate, and were expanded. Clones were screened by DNA sequencing and were differentiated to interrogate CDX2 protein expression by immunofluorescence. DNA sequencing primers sequences are: GCATCCTCCTGCTTCAGTCT and GCAGTTCTCAGCCCTCACTT. Control cells were treated the same way, but were electroporated with a GFP plasmid lacking a sgRNA.

### Generating doxycycline-inducible CDX2 expression in hESCs

The pInducer20 vector is an inducible Lentiviral vector system, which carries both rtTA3 and Neomycin-resistance genes under the UBC promoter as well as a cDNA of interest under the control of a tetracycline-responsive promoter (Meerbrey et al., 2011). pInducer plasmids are available from Addgene (44012). Generation of the pInducer-GFP inducible control cell line has been described elsewhere (Chen et al., 2014). The human *CDX2* clone HsCD00045643 was purchased from the Arizona State DNA Plasmid Repository (Dnasu.org), and was introduced into the pInducer20 plasmid using Gateway recombination cloning technology (ThermoFisher Scientific) following standard manufacture protocols. Lentiviral particles were generated at the University of Michigan Viral Vector Core, and inducible hESCs lines were generated as previously described (Tsai et al., 2016). Following antibiotic selection (G418), GFP or CDX2 gene expression was induced by adding 2 µg/mL doxycycline to the culture medium.

### Western Blotting

The cells at different differentiation stages were collected, homogenized and lysed in lysis buffer. Lysis buffer consisted of 50mM Tris pH 7.4, 150mM NaCl, 1% Triton X-100, 1.5mM MgCl_2_, 5mM EGTA, 1% glycerol and protease and phosphatase inhibitors (30mM Sodium Pyrophosphate (Na_4_P_2_O_7_), 50mM Sodium Fluoride (NaF), 0.1mM Sodium Orthovanadate (Na_3_O_4_V) and Complete, EDTA-free (Roche #11873580001). Cells were lysed for 30min at 4°C with rotation and cleared by centrifugation. Protein concentrations were determined by Bio-Rad protein assay (Bio-Rad #500-0006). Samples containing 20μg of protein lysate in laemmli sample buffer were separated on 15% SDS-PAGE gel and transferred (semi-dry) to nitrocellulose membrane. Membrane was probed for rabbit mAb CDX2 (D11D10) (Cell Signaling Technology #12306), then re-probed for mouse mAb β-actin (Sigma #A1978). HRP-conjugated rabbit or mouse secondary antibodies (Cell Signaling Technology #7074 or #7076) were used and the membrane was developed by using chemiluminescence.

### Purification of Intestinal epithelial cells from mice

Pregnant dams were sacrificed and dissected embryos were kept in ice cold PBS. Embryo tail tissue was used for genotyping using KAPA Mouse Genotyping Kits (Kapa Biosystems, KK7352). For E12.5 RNA-seq entire gut tubes (caudal of stomach), and for E16.5 ATAC-seq midguts (caudal stomach to rostral caecum) were dissected out of the body cavity. Intestinal tissues were treated with pre-warmed 0.25% Trypsin for 8-10mins at 37°C on a vortex station, neutralized with 10% FBS, and passed through a 70µm filter to obtain single cells. Single cells were incubated with PE-conjugated anti-CD326 (EpCam clone G8.8, eBiosciences, 12-5791-81) for ~30 minutes on ice. PE-stained cells were then incubated for ~30 minutes with magnetic conjugated anti-PE antibody (Miltenyi Biotec Anti-PE MicroBeads, 130-048-801). Subsequently, upon the availability of Anti-CD326 (Epcam) magnetic microbeads antibody (Miltenyi Biotec 130-105-958), cells were stained with the Anti-Epcam magnetic microbeads antibody for ~40mins minutes on ice. Stained cells were passed through a column (Miltenyi Biotec, MS Columns, 130-042-201) in a magnetic field to obtain magnetic antibody conjugated, EpCam positive epithelial cells. The purity of magnetic cell isolation was compared to FACS sorted EpCam positive cells and found to be comparable. Cells were dissolved in Trizol for RNA processing or used immediately for ATAC-seq.

### ChIP-seq

For embryonic CDX2 ChIP-seq (E13.5, E16.5, E17.5) midguts (caudal stomach to rostral caecum) were collected. Gut tubes were cut into small pieces, pooled, and fixed in 1% formaldehyde for 20-30mins at RT. Tissues were PBS washed twice and flash frozen for subsequent chromatin pulldown assays. Cell pellets were thawed on ice and dissolved in 3X volume of lysis buffer: 1%SDS; 10mM EDTA; 50mMTris-HCl pH8.1; Protease Inhibitor 1X (Complete inhibitors, Roche) or mammalian Protearrest (G biosciences). Cells were sonicated (~25-30mins) to shear chromatin in 200-600bp fragments in a Diagenode bioruptor. Chromatin was incubated overnight with CDX2 antibody (6µl, Bethyl A300-691A) conjugated to protein A/G beads (15µl each). Chromatin bound beads were washed 5X with RIPA wash buffer (50mM HEPES pH7.6; 1mM Edta; 0.7% Sodium deoxycholate; 1% NP-40; 0.5M LiCl) and rinsed once with TE buffer (10mM Tris + 0.1mM EDTA). Chromatin bound beads were re-suspended in reverse crosslinking buffer (1% SDS + 0.1M NaHCO3) and incubated at 65 oC for 6 hours to release ChIP DNA. DNA was column purified using a PCR purification column (QIAGEN) and quantified using Picogreen (Life Technologies). ChIP DNA was used to prepare ChIP-seq libraries using Rubicon Genomics ThruPLEX DNA-seq Kit (R400427/R400428/R40048), fragment size selected using Pippin Prep and sequenced on Illumina HiSeq (50 or 75-bp reads; single end; ~25M reads).

### ATAC-seq

25,000-50,000 isolated epithelial cells, from midguts (caudal stomach to rostral caecum, up to 2 embryos per sample) were used for ATAC-seq as described previously (Buenrostro et al., 2015) with minor modifications. Briefly, cells were centrifuged 500g for 5mins, and lysed in ice cold ATAC-lysis buffer (10 mM Tris-Cl, pH 7.4, 10 mM NaCl, 3 mM MgCl2, 0.1% (v/v) NP-40), followed by centrifugation at 500 g for 10 minutes at 4°C to isolate nuclear pellets, which were treated in a 50 μl reaction with Nextera Tn5 Transposase (Illumina, FC-121-1030) for 25-30 mins at 37°C. The transposed chromatin was purified with QIAquick PCR Purification Kit (QIAGEN) and PCR amplified with high-fidelity 2X PCR Master Mix (New England Biolabs) using an universal forward primer and unique reverse primers in a 50µL reaction for 5 cycles. After 5 cycles, 5µL of the reaction mix was amplified by qPCR for 20 cycles to determine the optimum number of additional cycles, based on 1/4 the maximum fluorescent intensity. The PCR amplified libraries were column purified, fragment size selected using Pippin Prep and sequenced on Illumina HiSeq (50/75-bp reads; single end; ~25M reads).

### Computational analysis

Raw Sequencing reads (fastq) were quality checked using fastQC (v0.11.3), aligned to mouse (mm9) or human (hg19) genomes using Tophat2 (v2.1.0, RNA-seq) or bowtie2 (v2.2.6, ChIP- and ATAC-seq) to generate bam files. For ChIP- and ATAC-seq, bam files were merged using samtools merge (0.1.19) for downstream analysis. Bigwigs were generated from bam files using deeptools bamCoverage (v2.4.2, (Ramirez et al., 2014), duplicate reads ignored, RPKM normalized and extended reads for Chip- and ATAC-seq) and visualized in Integrated Genomics Viewer (IGV, (Robinson et al., 2011).

RNA-seq: Cxb files were generated from BAM files using Cuffquant (v2.2.1, frag-bias-correct, multi-read-correct). Normalized expression count tables were generated from cxb files using Cuffnorm (v2.2.1, library-norm-method quartile). Differential gene expression tables were computed from cxb files using Cuffdiff (v2.2.1, –multi-read-correct –frag-bias-correct, –dispersion-method per-condition –library-norm-method quartile) and log2 fold change (FPKM+1) used for analysis.

For ChIP-seq and ATAC-seq: Peaks (bed file) were identified from aligned reads (bam) using MACS (1.4.1, Zhang et al., 2008). Two biological replicate ChIPs were performed for each condition, and replicates inspected by measuring Pearson correlation coefficients. Genes associated with ChIP peaks were identified using BETA minus (genes within 5kb of binding sites, using CTCF boundaries to filter peaks around a gene). Enriched Motifs at ChIP peaks were identified using HOMER findMotifsGenome.pl (v4.8.3, knownResults, (Heinz et al., 2010). Enriched ontologies were identified from genomic regions (bed file) using GREAT analysis (v3.0.0) using the Proximal- 5kb upstream and 1kb downstream and Distal 200kb setting (McLean et al., 2010) or DAVID (Huang da et al., 2009).

For comparing open chromatin regions (ATAC-seq) or differential CDX2 binding (ChIP-seq), bigwigs were generated (no normalization) and signal intensities were quantile normalized (v0.4.0, bin_size 50, log transformation) using Haystack (Pinello et al., 2018). *k* -means clustered heatmaps were generated from haystack normalized bigwigs using computeMatrix and plotHeatmap from deeptools package (v 2.4.2). Genomic regions (bed file) associated with desired k-mean clusters were extracted from bed files generated by PlotHeatmap, to obtain a selected bed files (genomic regions).

To identify condition-specific binding sites, MACS called peaks at P-value 10^-3^ for E13.5 and at P-value 10^-3^ and 10^-5^ for Adult were concatenated for each biological replicate, yielding a total of 33,834 genomic regions. E13.5 and Adult replicate bam files were merged respectively, and quantile normalized bigwigs were used for k-mean clustering (k=4) at these sites. Cluster2 was identified as Cdx2’s E13.5 enriched genomic targets (4,132) and cluster 3 as Cdx2’s adult enriched genomic targets (12,551 regions, genomic coordinates provided in Table S3). For human data, replicate bam files were merged for subsequent analysis. MACS peaks at P-value 10^-10^ were called on merged bam files of hindgut-specified endoderm (58,981 peaks) and Adult (19,752 peaks). *k*-means clustered heatmap was generated using quantile normalized bigwigs at Hindgut-specified endoderm (top cluster) and Adult sites (bottom cluster). Genes within a 5KB region of CDX2 summits were identified using BETA minus at hindgut-enriched and adult-enriched binding sites, and nearby genes were further used for evaluating gene ontologies (Table S1). For CDX2 ChIP-seq in naïve endoderm with doxycycline-induced CDX2, two biological replicates were performed and replicate bam files were merged for subsequent analysis. MACS peaks were called on merged bam files. Genomic sites bound by CDX2 in the doxycycline-induced condition were identified as overlapping sites of hindgut-specified endoderm using intersectBed peaks at P-value 10^-10^ (23,076 peaks). Condition-enriched sites were identified by subtracting lower stringency peaks of one condition (P-value 10^-3^) from higher stringency peaks of the second condition (P-value 10^-10^). This approach yielded Hindgut-only peaks (30,501 sites), not bound by CDX2 in the Tet-On condition, and Tet-On enriched peaks, which were bound upon CDX2 induction without differentiating the cells towards intestine (87,373 sites).

For ATAC-seq, two biological replicates from each developmental stage were performed, bam files were merged and bigwigs were quantile normalized to compare intensities of open chromatin signal. Embryonic and adult ChIP-seq data (Kazakevych et al., 2017) of the active chromatin marker (H3K27ac) was applied to Cdx2’s E13.5 and adult enriched genomic targets. Quantile normalized bigwig files of different replicates were merged by BigWigMerge. To evaluate whether CDX2 is required to maintain open chromatin regions where it typically binds, we compared ATAC-seq at E16.5 intestinal epithelium with CDX2 KO (*Shh-cre*) at E16.5. Two biological replicates were performed, bam files were merged and bigwigs were quantile normalized to compare intensities of open chromatin signal at regions indicated above.

### Statistical analysis

For statistical analysis associated with Figure 1, data are expressed as the median of each sample set. Each data point in the plots represents an independent biological sample. For organoid experiments, each independent biological sample is comprised of 3-5 organoids pooled together. All organoid experiments were conducted on at least 3 independent biological replicates. Unpaired t-tests, were carried out with GraphPad Prism 5.0 software. Each data point is presented, with the middle line representing the Mean, with error bars representing +/- SEM. In all figures, * = P<0.05. GSEA was run using the pre-ranked setting on all genes using FPKM + 1 values as described (Subramanian et al., 2005).

## Acknowledgements

Research reported in this publication was supported by National Institute of Diabetes and Digestive and Kidney Disorders (NIDDK) and National Institute of Allergy and Infectious Diseases (NIAID) of the National Institutes of Health under grant number U01 DK103141 (MPV and JRS). The content is solely the responsibility of the authors and does not necessarily represent the official views of the National Institutes of Health. This work was also supported by the Human Genetics Institute of New Jersey; NIH grants R01 CA190558 (MPV), R01 DK082889 (RAS); the Biospecimen Repository Service Shared Resource and Sequencing Facility of the Rutgers Cancer Institute of New Jersey (P30CA072720); and the University of Michigan Center for Gastrointestinal Research (UMCGR) (NIDDK 5P30DK034933).

## Author Contributions

Conceptualization, MPV and JRS; Investigation, NK, YHT, LC, SH, KKB; Analysis, NK, LC, AZ, KKB, MS; Supervision, MPV, JRS, JX, RAS; Writing – Original Draft, NK and MPV; Writing – Reviewing and Editing, NK, MPV, JX, RAS, JRS.

## Supplemental Figures

**Figure S1.**
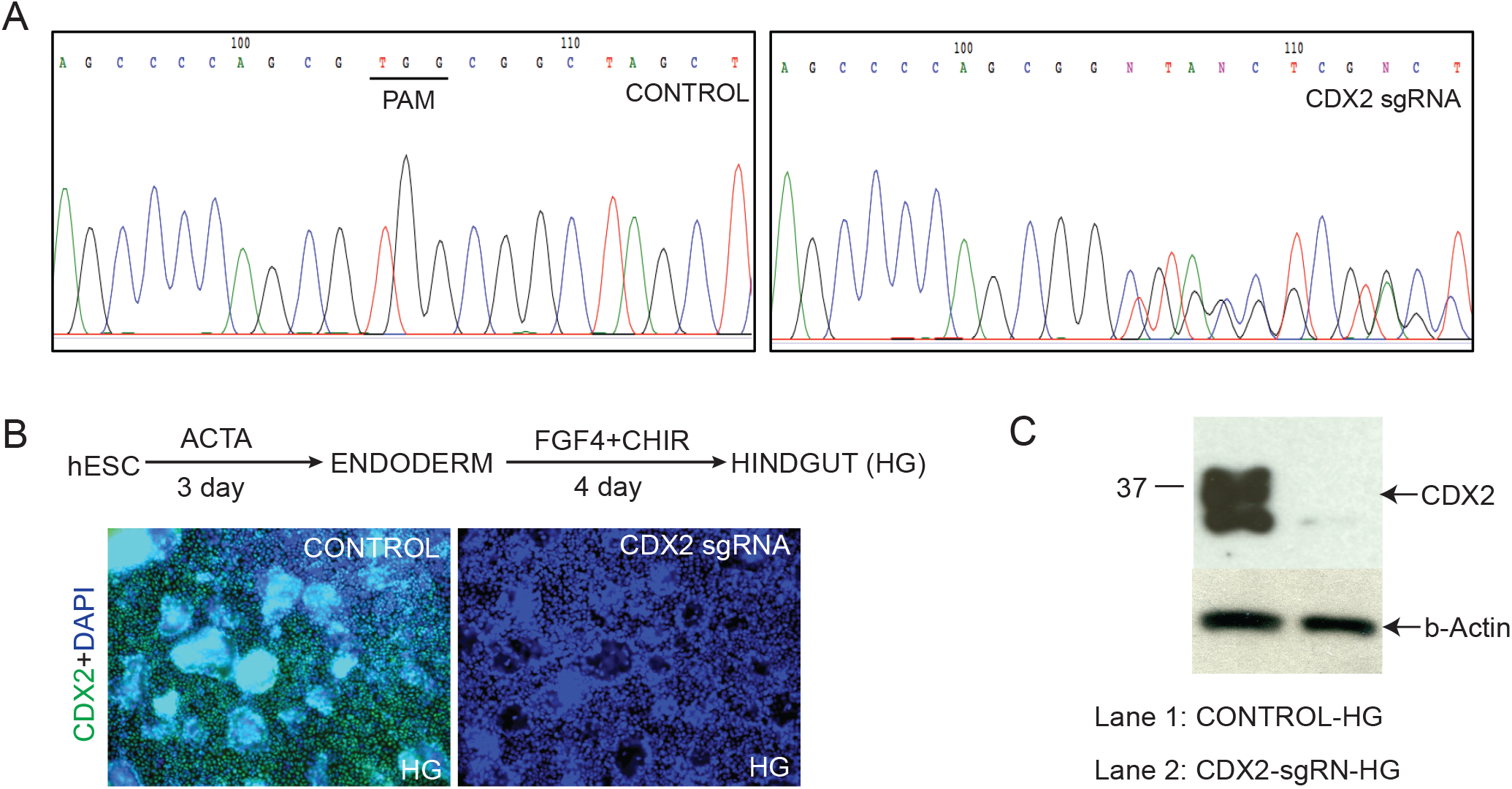
The *CDX2* coding region is disrupted by Crispr targeting (A), resulting in loss of CDX2 protein expression as detected via immunostaining of cells following hindgut-directed differentiation (B) or by immunoblotting (C).

**Figure S2.**
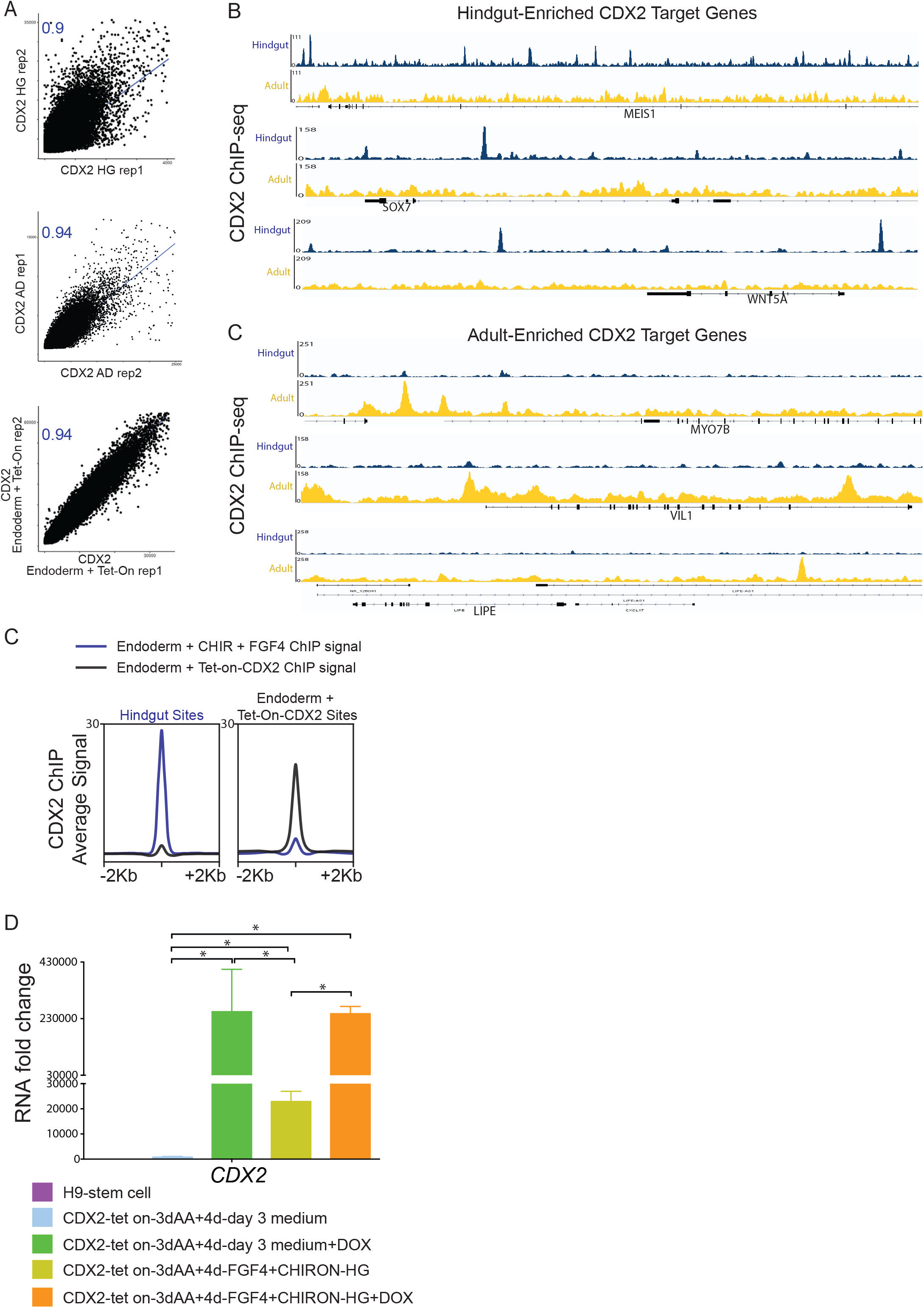
CDX2 has distinct transcriptional targets in the developing hindgut versus the adult intestine. A) Pearson Correlation plots indicate consistency between ChIP-seq replicates. (B) Representative examples of CDX2 binding to distinct targets in hindgut versus adult intestinal epithelium. C) Average Signal for CDX2 binding at normal hindgut sites, versus sites bound by CDX2 only when induced via doxycycline (Ectopic CDX2 sites, Figure 2F). D, *CDX2* transcript levels are robustly induced upon Dox-treatment in the CDX2-tet on cell line, as indicated by qRT-PCR.

**Figure S3.**
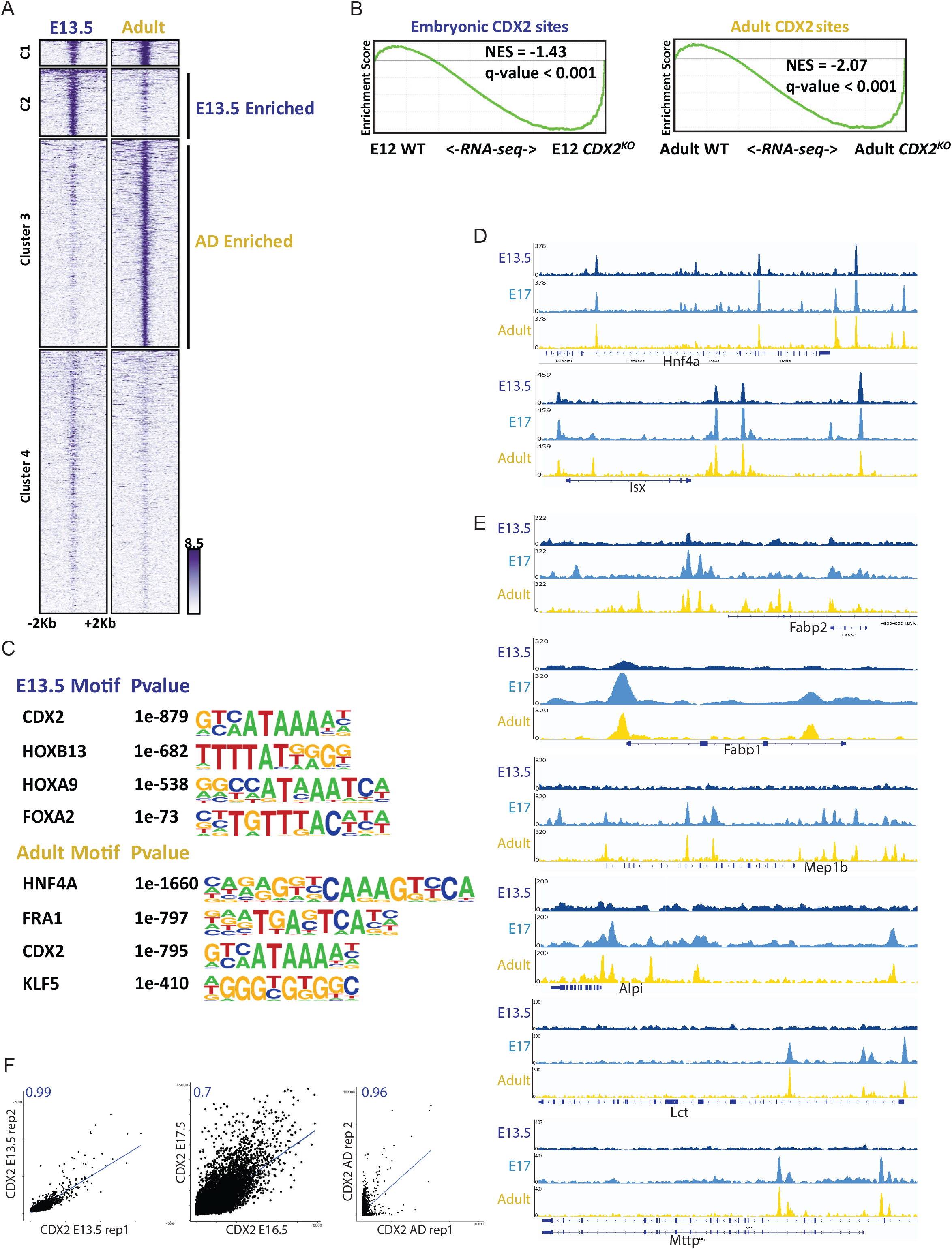
Temporal-specific CDX2 binding is conserved between mice and humans, and impacts intestinal gene expression. A) *k*-means clustering facilitated identification of sites occupied by CDX2 robustly in the intestinal epithelium at either E13.5 or adult stages. B) Genes within 5kb of a condition-specific CDX2 binding site were dependent upon CDX2 for expression, as these genes tend to decrease upon knockout of CDX2 in the embryo (*Shh-Cre; Cdx2^f/f^*) or adult (*Villin-Cre^ERT2^; Cdx2^f/f^*), as measured by RNA-seq. C) Distinct classes of DNA-binding motifs are enriched at CDX2-binding regions unique to the embryo or adult stages, suggesting that CDX2 has distinct partner factors at these developmental stages. D) Examples of persistent, or E) developmental stage-specific, CDX2 binding in the mouse. F) Pearson Correlation plots indicate consistency between ChIP-seq replicates.

**Figure S4.**
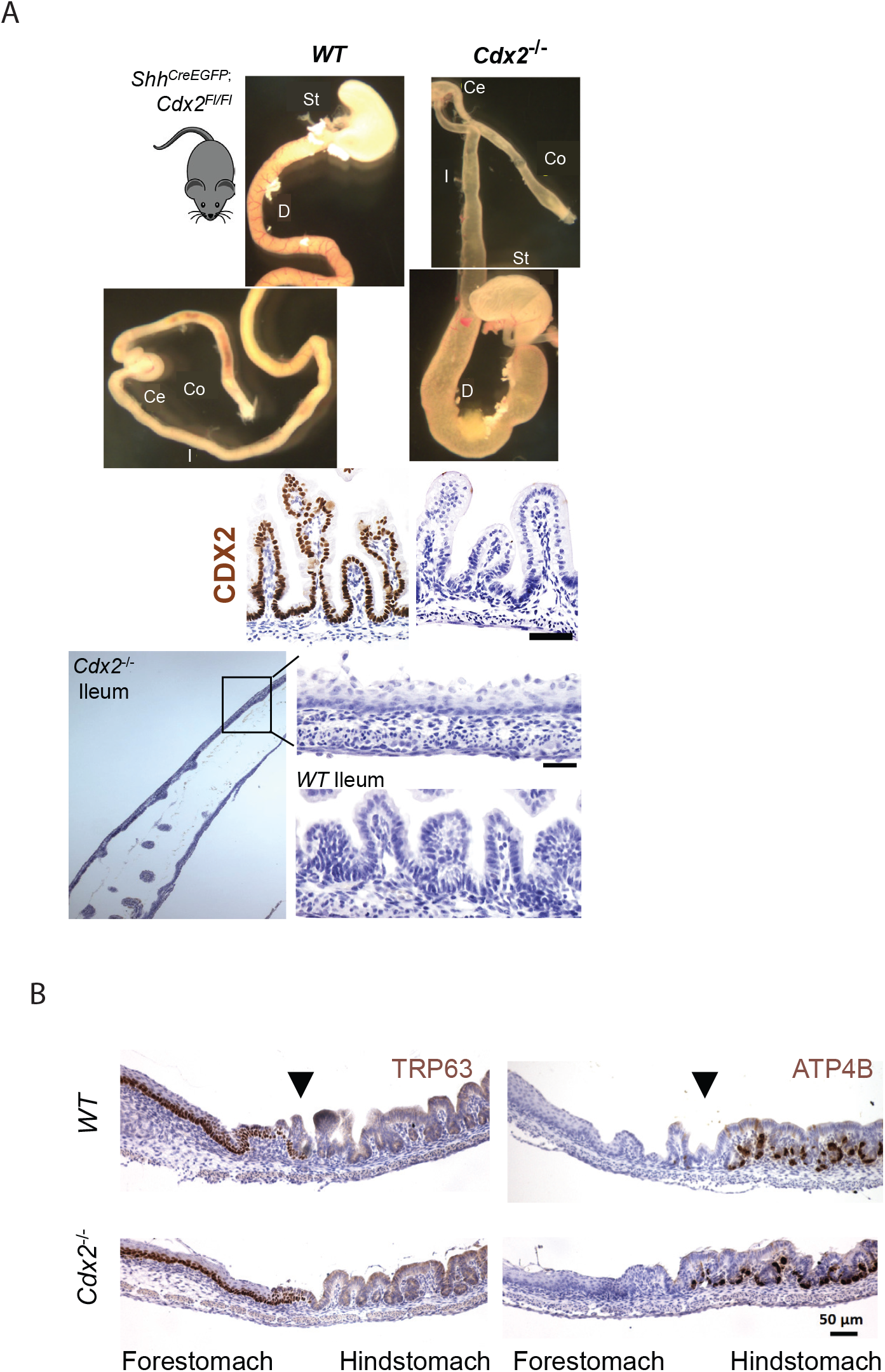
(A) At the top, whole mount images depicting the *Shh^CreEGFP^;Cdx2^f/f^* phenotype at E18.5. Note the distended intestinal lumen. Below, CDX2 immunostain indicates efficient deletion of *Cdx2*. A closer look at the ileum of the mutants reveals a keratinized squamous appearance in lieu of the simple columnar epithelium present in the control. (B) Control staining of stomachs from *Shh^CreEGFP^;Cdx2^f/f^*mutants and controls shows the expected pattern of immunoreactivity for TRP63 in the squamous forestomach, and ATP4B in the glandular hindstomach. These stains were done in parallel, and as controls, for those shown in Figure 4.

**Figure S5.**
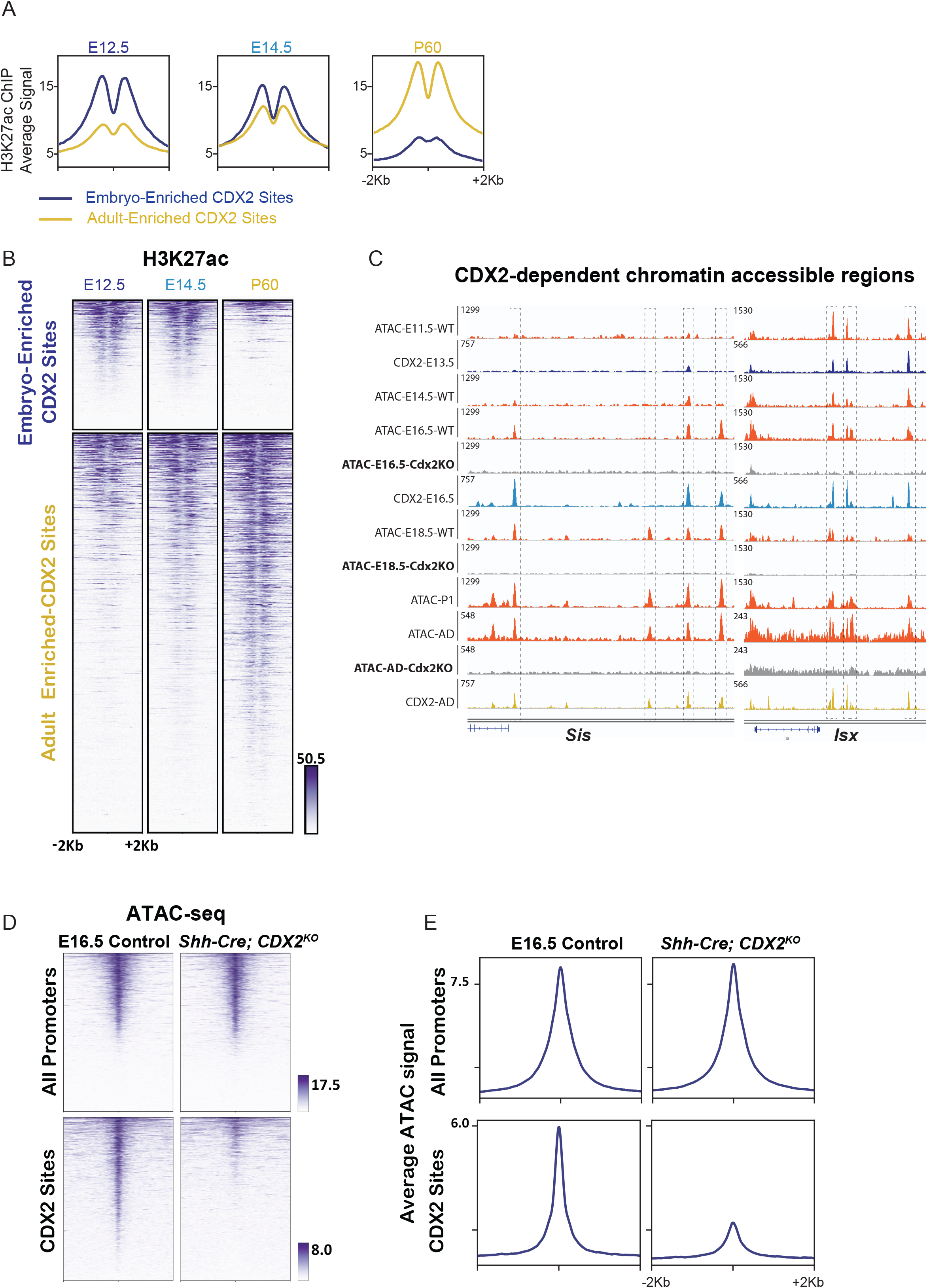
(A-B) Composite plots and heatmaps demonstrating the active chromatin marker, H3K27ac, is enriched at CDX2-binding sites in a stage-specific manner. H3K27ac ChIP-seq data are from (Kazakevych et al., 2017). C) Example data traces of CDX2-binding at accessible chromatin regions over developmental time (ATAC-seq), and their dependence upon CDX2 expression to remain accessible upon CDX2 knockout using the *Shh-Cre* or *Villin-Cre^ERT2^* (gray tracks). (D-E) Substantial loss of accessible chromatin is observed at CDX2-bound genomic regions in E16.5 *Shh-Cre*; *Cdx2^f/f^* embryos, whereas promoter regions are relatively unaffected.

## Supplemental Tables

Table S1. Genome coordinates for CDX2-binding data from ChIP-seq performed in human hindgut (Hindgut-enriched sites) and Adult (Adult-enriched sites). Additionally, the results of HOMER motif-calling analysis on these sites enriched more specifically in the hindgut or adult are reported. Finally, the results of GO term enrichment using DAVID analysis for genes in proximity to these binding regions are reported. These data correspond to findings displayed in Figure 2.

Table S2. CDX2 binding data from ChIP-seq performed in Hindgut (Activin, WNT/FGF treatment protocol) versus Endoderm treated with Doxycycline (Activin, Doxycycline-CDX2). Binding coordinates of CDX2 sites found in both conditions (OverlapHindgutTetOn), sites more robustly bound by CDX2 in hindgut (HindgutEnriched), or sites more robustly bound by CDX2 when ectopically expressed in endoderm without the hindgut-inducing WNT/FGF-treatment (TetOnEnriched) are listed. Additionally, the results of motif-calling analysis of the sites enriched more specifically in the Tet-On or WNT/FGF treated condition are reported. These data correspond to findings displayed in Figure 2.

Table S3. Genome coordinates for CDX2-binding data from ChIP-seq performed in purified mouse intestinal epithelium at E13.5 (Embryo-enriched sites) and Adult (Adult-enriched sites). Additionally, the results of motif-calling analysis on these sites enriched more specifically in the mouse embryo or adult are reported. Finally, the results of GO term enrichment using GREAT analysis for genes in proximity to these binding regions are reported. These data correspond to the findings in Figure 3.

